# Scaling up naturalistic narrative research: A reproducible large language model-based tool for evaluating contextual information gain

**DOI:** 10.64898/2026.11.14.766702

**Authors:** Rohin A. Palsule, Aidan Goeschel, Jungsun Yoo, Janice Chen, Aaron M. Bornstein, Ari Khoudary

## Abstract

Narratives are a universal part of natural human experience. As experimental stimuli, narratives allow scientists to study complex mental processes with direct relevance to everyday experience. These analyses typically require manual labeling of key event features. Such characterization of narrative stimuli are extremely labor-intensive and prone to subjectivity. Automated tools could reduce these difficulties and enable scaling to longer, varied stimuli by exposing the exact quantities computed. We address this by introducing a metric that uses open-weight LLMs to compute *contextual information gain*. Our metric correlates with human ratings of event importance and causal centrality, critically distinguishing the latter from semantic centrality, thus overcoming limitations of existing automated tools. The metric similarly captures shared coactivation in the default mode network, a network implicated in narrative comprehension. Taken together, these results validate an automated, empirically-grounded tool that researchers can use to identify behaviorally- and neurally-relevant computational quantities in naturalistic stimuli.

**Research Transparency Statement:** Conflicts of interest: None. Funding: This work was supported by NIH R01AG088306 and R01NS119468. Artificial intelligence: Artificial intelligence use was limited to the visual clarity of figures and supporting plots. No artificial intelligence was used for the generation of text in the paper. Ethics: All experimental procedures and analyses were in accordance with protocols approved by the Institutional Review Boards (IRB) of Johns Hopkins. Preregistration: None of the reported analyses were preregistered, all subject data was adapted from open source studies and cited accordingly.

## 1. Introduction

Narrative stimuli—spoken stories, animated videos, and television shows/movies—are powerful tools for studying naturalistic cognition (Lee et al., 2020). A stimulus can be called “narratively structured” if it consists of a series of events that occur in temporal succession of each other, and especially so if there are causal links among events that are not temporally contiguous (Chen & Bornstein, 2024). These stimuli are thought to resemble the complexity, contents, and statistics of real-world experience, thus making them more ecologically valid for the study of information processing. For example, narrative stimuli have accelerated our understanding of the neurocognitive mechanisms driving event perception, a process that is central for organizing experiences in episodic memory (Zacks et al., 2020).

A recent proposal suggests that humans’ ability to efficiently integrate temporally discontiguous information is the result of utilizing narratively structured schemas to compress the enormous *state space* of the natural world into tractable, discrete, interconnected representations (Chen and Bornstein, 2024). These schemas allow humans to generate both retrospective causal relationships and prospective predictive probabilities of event representations as they unfold. The causal structure of these narrative schemas is especially relevant given the mounting evidence from narrative-based neuroscience research demonstrating that human memory systems are distinctly sensitive to causal information compared to temporal ordering (Antony et al., 2024; Cohn-Sheehy et al., 2021; Lee & Chen, 2022; Lee et al., 2020; Zadbood et al., 2022).

Naturalistic neuroscience research allows for the online evaluation of neural regions involved in processing the complex structure of these stimuli. Crucially, the default mode network (DMN), neural regions originally implicated in “offline” mind-wandering and contrasted against attentional processes (Buckner et al., 2008; Raichle, et al., 2001, 2015), has been highlighted as responsible for long-term processing and evaluation of narrative events (Ritchey et al., 2015; Barnett et al., 2020; Baldassano et al., 2018; Chen et al., 2017; Simony et al., 2016) as well as attentional engagement (Song et al., 2021; Estermen et al., 2013) regardless of temporal ordering.

Methodological advancements in neuroimaging research allow for the evaluation of shared neural coactivation while processing narrative content, and additionally allow for the comparison of these shared signals with stimuli-related metrics. Specifically, researchers have connected time-varying cognitive measures with inter-subject functional correlations (ISFC) of brain responses to find that DMN activity fluctuates according to event-by-event surprisal, along with comparing measures of importance, emotional valence, and vividness with this network coactivation (Brandman et al., 2021).

Critically, existing automated methods are limited to assessing the semantic similarity of events (Lee & Chen, 2022), which only partly captures the richness of psychological experiences empirically linked to narrative processing. Previous research has shown overlapping effects of semantic similarity and manually identified causal similarity (Lee & Chen, 2022), emphasizing the need for distinguishable automated metrics capturing cognitive measures. The other manual measures require independent raters to laboriously generate them, which is both time consuming and lacks any automation for inter-study meta-analyses across different naturalistic stimuli. For example, a dataset discussed in this article required 8-12 independent raters per measure during data acquisition. Additionally, such ratings are prone to individual and group subjectivity that could differ from the intended interpretation of each metric.

Large language models (LLMs) may serve as a useful tool for automating the labeling of event-level properties related to cognitive processes. The massive scale of data on which LLMs are trained makes them reasonable proxies for ground-truth frequencies about token co-occurrences, which could plausibly extend to higher-level properties such as schemas. The variability in subjective interpretation of these properties in independent human raters, usually addressed by averaging, can be mitigated by using reproducible software-based pipelines. Further, the attention mechanism in transformers is uniquely capable of identifying token relations across large windows of textual “evidence,” facilitating connections between events that are conceptually related but temporally discontiguous. These token relations allow for human-like conceptual representations of internal representations (Du et al., 2025), a relationship also displayed with traditional word embeddings (Caliskan et al., 2017). Taken together, these properties of contemporary LLMs make them suitable candidates for investigating how complex, high-dimensional prior knowledge might shape real-time encoding of continuously unfolding information.

We begin by adapting an existing LLM-based metric, *Sequentiality*, that was originally interpreted as representing “narrative flow” (Sap et al., 2022). While the original implementation was based on GPT-2, here we re-implement using an ensemble of open-weight models. We first validate that our implementation accurately reproduces the original findings. Then, we show that the evidence produced by the model is useful for reproducing existing labelings of naturalistic narrative stimuli when the event probabilities are re-computed as a measure of *contextual information gain*, conditional on an experimenter-defined “topic” that represents the subject’s goals or awareness of the eventual outcome. To validate this new tool, we re-analyzed several previously published datasets demonstrating behavioral or neural responses to narratively-structured stimuli. Specifically, we evaluated Sequentiality’s sensitivity to (1) the importance and causal structure of narratives, (2) shifts in prior expectation during narrative-twists, and (3) event boundaries. We then examined both univariate activation modulated by Sequentiality as well as intersubject functional correlations to model how DMN activity maps onto Sequentiality relative to other manual cognitive measures.

## 2. Methods

### 2.1 Original Sequentiality formalization

We build upon an existing metric, *Sequentiality*, that was created to quantify the narrative “flow” of output events in text-based narrative stimuli (Sap et al., 2020, 2022). This metric derives likelihood values for each event (or sentence) in a narrative using token probabilities obtained via LLMs. Specifically, the likelihood of a particular sentence is defined by two generating models: one based on a user-defined topic τ only, and one based jointly on the topic and preceding h sentences. The original Sequentiality metric thus defined the “flow” *c* of a narrative text as the difference in the likelihood of sentence *s*_*i*_ between these two generating models:

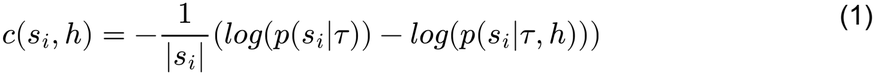

where *s*_*i*_ is sentence *i, τ* is the topic that sets the situation model for the story, and *h* is the history length defining number of previous sentences *s*_*i−h*:*i*−1_ that are used to establish a local context. The values of *h* and τ are defined according to experimenter discretion and thus can facilitate different applications of the metric. Details on how we define these values for our analyses are contained in Section 2.5.

The first term inside the parentheses in Equation 1 corresponds to the likelihood of sentence *s*_*i*_ conditioned only on topic *τ*, and the second term corresponds to the likelihood of sentence *s*_*i*_ conditioned jointly on topic *τ* and previous context *h*. Essentially, this measure quantifies how much the previous sentences in the context *reduced* the surprise of observing sentence *s*_*i*_. A positive Sequentiality value for a sentence *s*_*i*_ thus indicates that the previous *h* sentences made the probability of sentence *s*_*i*_ *greater* than expected by the topic alone, whereas negative values indicate that the previous *h* sentences made the probability of *s*_*i*_ *lower* than what was expected based on the topic (Sap et al., 2022). Finally, the metric is normalized by sentence *s*_*i*_ to control for differences in sentence lengths within the narrative.

### 2.2 Formal equivalence to contextual information gain

Although Sequentiality was originally developed to formalize a notion of narrative “flow,” here we show that it can be equivalently re-expressed as a metric interpretable as contextual information gain. We begin by adding the topic term *τ* as an explicit parameter, exposing the interpretation of the measure as conditional on an explicit prior outcome or goal. We then combine the log terms in Equation 1:

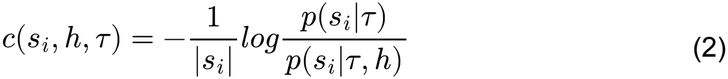

To better conceptualize the meaning behind this ratio, we can expand the terms in this equation using Bayes’ Rule:

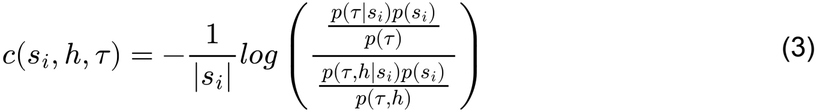

and then simplify to obtain:

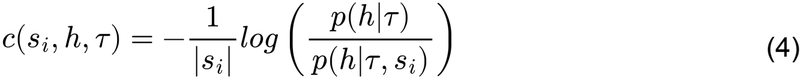

This final simplification represents how much information each new observation *s*_*i*_ carries about the context established in history *h* that is *not* already predicted through the topic *τ* alone. It is precisely this quantity that we term *contextual information gain*.

### 2.3 Open-source implementation

A computational metric quantifying the dynamic flow of information in a narrative is a powerful tool for scaling up naturalistic neuroscience research. However, its applications outside of the original study are limited by the proprietary nature and computational demands of the GPT2 implementation. To address this gap, we develop and validate an open-source version of the Sequentiality metric. Importantly, our implementation automatically generates likelihoods from five different LLMs with varying architectures, training sets, and fine-tuning protocols, allowing for more robust conditional probability estimation and/or investigation into systematic variations in Sequentiality as a function of model provider. Specifically, our implementation utilizes likelihoods from Sakana TinySwallow 1.5B, GPT2-XL, OLMo-2-13B, Llama-3.1-8B, and Qwen2.5-7B. The Python code used to implement this open-source Sequentiality metric is available for public use on Github and can be readily adapted to incorporate new open-weight models as desired.

To calculate Sequentiality, we first tokenize the entire story using the same alphabet as each LLM in the ensemble. To obtain topic-only sentence likelihoods, *p*(*s*_*i*_|*τ*), we append a preamble to our input text that tells the LLM to adjust its probabilities according to a user-defined topic *τ*. Then, for each token *t* in sentence *s*_*i*_, we pass in all tokens before *t* in *s*_*i*_ and extract the distribution for the predicted next token. We can search through this distribution to find the log-probability of token *t* given the previously inputted tokens. These log-probabilities are summed to generate the topic-driven likelihood, *p*(*s*_*i*_|*τ*).

To calculate the context-driven sentence likelihood, *p*(*s*_*i*_|*τ, h*), we use the same preamble to establish a topic and the same mechanism of autoregressive distribution generation. However, rather than inputting only *s*_*i*_, we input the previous *h* sentences as well, where *h* is a hyperparameter that controls the history length of the model. The Sequentiality value for a single sentence *c*(*s*_*i*_, *h, τ*) is then computed as specified in Equation 1: we compute the difference between topic and context likelihoods and then normalize by the number of tokens in the sentence.

Our method thus generates *h* * *M* Sequentiality values for each sentence in a piece of written text, where *h* is the history length and *M* is the number of models in the ensemble. A primary strength of this method is the ability to quantify and represent uncertainty about Sequentiality values across model providers for a single sentence—or narrative—which was impossible to achieve with the previous approach. Further, as we show in Figure 1B, averaging across five open-weight LLMs generates Sequentiality values that are nearly identical to those obtained from the now-proprietary GPT-3. This is notable for two reasons. First, the open-source nature allows for broader and more transparent application of the Sequentiality metric. Second, we make use of models that are orders of magnitude smaller than originally used, making the computation of Sequentiality more tractable for researchers without access to massive computational resources, and which makes the tool more readily scalable to larger corpora with more diverse or extended stimuli.

**Figure 1:**
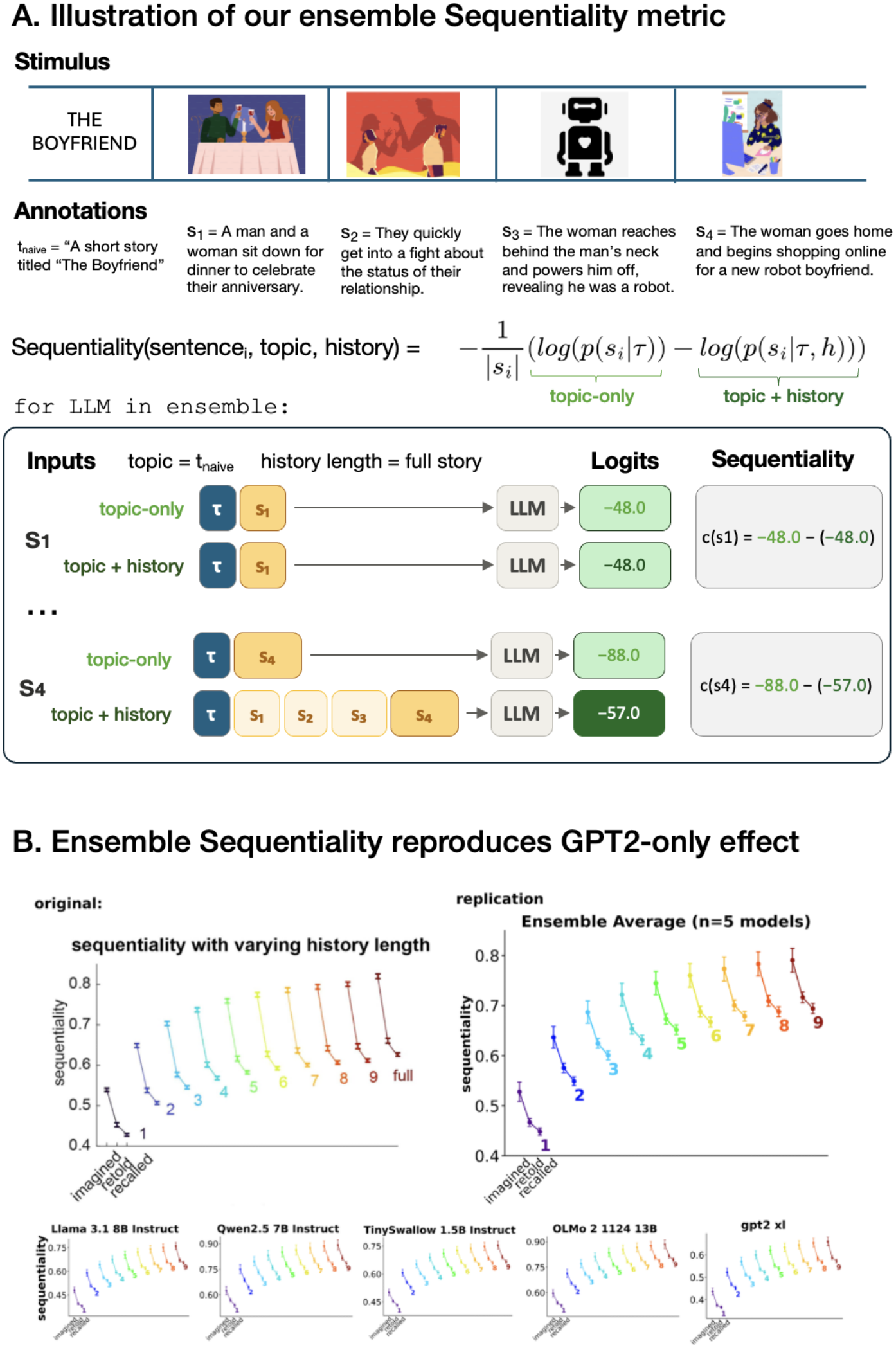
Open-source ensemble Sequentiality metric. (A) The Sequentiality metric receives inputs *τ*, or a user-defined sentence outlining the topic of the story, *h*, or a fixed number of sentences preceding the current event it uses as history, and *s*_*i*_, or the current sentence the metric is computing. The metric then uses an ensemble of LLMs (Sakana TinySwallow 1.5B, GPT2-XL, OLMo-2-13B, Llama-3.1-8B, and Qwen2.5-7B) to tokenize each sentence and generate the cumulative log probabilities when a given sentence is conditioned only on *τ*, and additionally when conditioned on *τ* and *h*. Then, the log ratio of these two values is standardized and output as the Sequentiality value for a given sentence. (B) Our ensemble-based implementation replicates the main finding reported by Sap et al. (2022).

### 2.4 Implementation validation

To confirm that our implementation accurately computes Sequentiality we used the Hippocorpus database to replicate the main findings reported by Sap et al. (2022). Hippocorpus consists of 6,854 short, written stories about memorable life experiences. Human subjects were asked to recall an autobiographical event that happened to them recently, retell this story 3-6 months later, or create an imagined short story. For each story, participants also generated a short phrase describing what happens in the story. More details about the dataset can be found in Sap et al. (2020), and the dataset itself can be found on HuggingFace (https://huggingface.co/datasets/allenai/hippocorpus).

We computed our ensemble Sequentiality metric for each sentence in each story in the Hippocorpus dataset at history lengths ranging from 1 to the full length of the story. Within each history length, we computed the total Sequentiality of a story as the mean of each sentence’s Sequentiality as generated by each model. The total Sequentiality value at each level of history length is then computed as the mean across model-specific story-level Sequentiality values. Following the approach of Sap et al (2022), Figure 1B displays Sequentiality values only for topics that contain stories generated in each of the three conditions (recalled, retold, imagined; n=984 total stories). This approach controls for variability in Sequentiality due to topic, thus allowing for more direct comparisons of the effect of story generation type on Sequentiality.

As shown in the top panel of Figure 1B, our implementation (right) almost perfectly replicates the original findings (left). A linear regression on Sequentiality values confirmed that we replicated the main effect of story type on Sequentiality. Both recalled (*β* = −0.091, *p* < 0.001) and retold (*β* = −0.069, *p* = 0.001) stories had significantly lower Sequentiality values than imagined stories. Additionally, history length monotonically increased Sequentiality across all story types (*β* = 0.027, *p* < 0.001). Further, the bottom panel of Figure 1B shows that each model in our ensemble independently reproduced the original Sequentiality findings qualitatively, while exhibiting a moderate degree of variability in the precise Sequentiality values computed (note the different y-axes across figures). Importantly, the top panel of Figure 1B shows that averaging across these individual estimates puts our ensemble metric back onto the same quantitative scale as the original Sequentiality values.

### 2.5 Hyperparameter settings: history length and topic

In order to generate Sequentiality values, both a history length *h* and topic *τ* must be defined. Whereas the original study manipulated history length and used subject-generated topics, our analyses use a fixed value of history length and experimenter-generated topics.

#### 2.5.1 History length

We fix history length *h* as the total number of sentences comprising an annotated narrative. As shown in Figures 1 and S1, increasing history length increases the magnitude of Sequentiality values for the Hippocorpus dataset. Thus, we chose to fix history length to a value that permits the entire narrative to fit into the context window.

#### 2.5.2 Topic

For all analyses except those involving foreknowledge of narrative twists (Section 4.3), we set the topic as “A short story about ‘*Movie title*’”. This type of topic aptly represents the amount of information observers have about the events of a narrative prior to observing its contents. For analyses capturing the effect of having knowledge of the narrative’s outcome, we constructed topics that concisely conveyed the key pieces of information comprising the narrative (e.g., “A short story about a robot woman who shuts off her robot boyfriend”). The full list of these “outcome-aware” topics is provided in Table S3.

### 2.6 Input standardization

Our analyses make use of two types of inputs to the Sequentiality metric: human-generated annotations describing the contents of audiovisual movies and computer-generated transcripts of auditory-only narratives. We developed a preprocessing protocol that standardizes both types of inputs in order to minimize surface-level syntactical differences that may generate spurious Sequentiality estimates. Specifically, we ensured that all inputs adhered to the following criteria: subject-object-verb word ordering, active voice, neutral tone, and no dialogue.

For computer-generated transcripts of auditory narratives, we used the NeMo Forced Aligner (Rastorgueva et al., 2023) to create time-aligned transcripts, and then used Claude Opus 4.5 to perform the standardization preprocessing. The specific prompts used are provided in the Supplemental Materials. Human-generated annotations were initially provided by a sample of trained raters (Section 2.7.1) and then manually standardized by author R.A.P. Importantly, the standardization process did not involve judgments about event boundaries, stimulus properties, etc. that were not already present in the original annotations. Accordingly, future work can develop robust pipelines for annotation standardization, perhaps by utilizing vision-language multimodal models (VLMMs) that can jointly process visual and semantic properties of audiovisual narrative stimuli.

### 2.7 Human subjects datasets

To demonstrate the utility of the Sequentiality metric, we apply it to three existing datasets that each utilized different types of narrative stimuli and cognitive tasks. Relevant details about each dataset are provided below, and further details can be found in the original studies that generated the data (Lee & Chen, 2022; Sava-Segal et al., 2026; Michelmann et al., 2023).

#### 2.7.1 FilmFestival Dataset (Lee & Chen 2022)

Lee & Chen (2022) obtained behavioral ratings and neural activity measured via functional magnetic resonance imaging (fMRI) while human subjects (12 female, 9 male; age 20-33, mean 26.6 years) watched a series of short films (n=10 independent films, average duration = 4.54 minutes). The order of movies was fixed across participants and fMRI data were acquired in two blocks, each of which contained five of the 10 total movies in the stimulus set. Six subjects were excluded due to excessive head motion in both the original study and the analyses reported here. A separate sample of trained human subjects (n=4) generated timestamped annotations for experimenter-defined scenes/events comprising each movie in the stimulus set.

##### Event Semantic Centrality

The semantic centrality of events in each movie were computed automatically using Google’s Universal Sentence Encoder (USE). Each event annotation generated by each of the annotators was represented as a 512-dimensional vector in USE space. After verifying inter-rater similarity (cross-annotator cosine similarity = .78), embedding vectors were averaged across raters. These embeddings were then used to construct a narrative network for each movie, where each event corresponded to a node and edge weights were defined as the cosine similarity between USE vectors. The semantic centrality of each event was then calculated as its normalized node degree.

##### Event Causal Centrality

An independent sample of n=18 human subjects made binary judgments about causal relationships between pairs of scenes in each movie. Each subject rated a subset of the total stimulus set, such that each film was evaluated by 12-13 raters. Causal judgments were permitted to be bidirectional, such that a rating of 1 can indicate that scene X caused scene Y or vice versa. These ratings were then used to construct causal networks, where each node corresponds to a scene and the edge weights are defined as the proportion of raters who identified a pair of scenes as causally related. The causal centrality of an event was again calculated as normalized node degree where edges were the normalized cause-effect likelihood in a bidirectional network.

##### Event Importance Ratings

Another independent sample of n=8 human subjects viewed all stimuli outside of the scanner to provide ratings of event importance. Half of the subjects made retrospective judgments of importance after watching each film to completion, and half made “online” judgments when prompted at the end of each scene. Ratings ranged from 1 (not important at all) to 10 (very important). Importance ratings were positively correlated with each other both between the two rating groups (r(202) = .67, p < .001, 95% CI = [.58, .74]) as well as within a rating group (r_online_=.65, r_post-view_=.55). Given the degree of similarity between these two types of ratings, we averaged across them to increase the reliability of the analyses that follow.

#### 2.7.2 Repeated Listening Dataset (Sava-Segal et al., 2026)

Sava-Segal et al. (2026) obtained behavioral ratings and fMRI signals while human subjects (n=36; 11 male, 25 female; age 18-33, median 20 years) underwent two consecutive listens of the same 8-minute auditory narrative (“The Dark End of the Mall”). Critically, the narrative contained a plot twist halfway through that causes participants to reinterpret the actions of characters within the narrative. During both the first and second exposures, participants rated how favorably they viewed one of the characters on a scale of -3 (very negative) to 3 (very positive). Specifically, they were asked during their first viewing: *“Overall, how much do you like Lucy?”* and during their second viewing: *“For your 2nd story, you have been assigned to listen to the same story again and complete the same prompt. For this 2nd listen of the same story, consider how your impression has changed. Because you have already listened to the story, we expect that your impressions of Lucy are different than your 1st listen. Given what you know about this story, what is your impression of Lucy now? Please use the slider to continuously rate your impression.”* Participants made these ratings continually in the fMRI scanner during narrative viewing.

#### 2.7.3 Event Segmentation Dataset (Michelmann et al., 2023)

Michelmann et al. (2023) obtained behavioral ratings of event boundaries as perceived across three independent auditory narratives (“Monkey”, “Pieman”, and “Tunnel”). For Pieman, participants listened to the audio narrative twice over and were asked to segment the 7.5 minute long narrative each time (n = 205, no gender and age information available in the original manuscript, further details can be found in Michelmann et al., 2021). Due to the length of Monkey in the Middle (30 minutes), groups of participants segmented different parts of the story while listening to it. Per Michelmann et al., “All participants listened to the full story but only segmented their respective section; a total of 30 participants segmented the first quarter, 35 participants segmented the second quarter, 31 participants segmented the third quarter, and 42 participants segmented the final quarter of the story (Kumar et al., 2022).” Human-generated event boundaries were collected from 10 participants for Tunnel Under the World (25.6 minutes) while listening to the narrative. Response vectors were derived at millisecond resolution, where each entry in the vector was 1 whenever there was a response within one second, and 0 otherwise. Then, these vectors were averaged across participants and smoothed with a Gaussian kernel width of 2 s in “Pieman”, and a different kernel for the other two stories that had a standard deviation of 0.2 s, leading to an interval of roughly 1 s. From there, consensus event boundaries were defined as peaks in this distribution of event boundary ratings. After transcribing and standardizing transcripts for each story, we created sentence-level binary masks indicating the presence or absence of an event boundary in that sentence.

### 2.8 fMRI analyses on FilmFestival data (Lee and Chen, 2022)

We preprocessed the raw fMRI data acquired by Lee and Chen (2022) using fMRIPrep 23.0.0 with standard settings (Esteban, Markiewicz, et al. (2019) RRID:SCR_016216). Functional images were motion corrected, skull-stripped, and distortion corrected with two B0 nonuniformity map acquisitions. Corrected images were then resampled to MNI152NLin2009cAsym space at the original functional resolution. Further details about original data acquisition and the specific preprocessing pipeline we used can be found in the Supplemental Materials, and additional preprocessing is explained according to the functional analysis.

#### 2.8.1 Univariate Activation Analysis

We conducted univariate analyses to investigate the relationship between Sequentiality and the fMRI blood-oxygen-level-dependent (BOLD) signal across participants. After obtaining each participant’s voxelwise activation across all movies, we conducted a linear regression of z-scored Sequentiality onto cortical activity, including as predictors nuisance regressors and a term accounting for the hemodynamic response delay. Nuisance regressors included the six motion parameters along with their temporal derivatives, squares, and squared derivatives, as well as the white matter and cerebral spinal fluid signal regressors. We applied a high pass filter of 1/128 Hz and spatially smoothed the data with a Gaussian kernel of full width at half maximum (FWHM) of 6 mm. Participant-level parameter estimate maps were then thresholded using the false discovery rate (FDR) correction across all voxels at q < 0.05 to account for multiple comparisons.

#### 2.8.2 Dynamic Intersubject Functional Correlations (dISFC)

We utilized intersubject function correlations (ISFC) to measure across-participant activity in regions spanning different cortical networks (Simony et al., 2016). The ROIs used in this analysis were adopted from Yeo et al. (2011) and include the 114 parcel atlas spanning 7 functional networks that were calculated using the resting-state functional neural data of 1,000 healthy adults. The 114 parcels along with the functional data all remained in MNI space for these analyses.

Similar to previous studies using a dynamic, sliding temporal window for functional connectivity (Brandman et al., 2021; Song et al., 2021), we utilize a 15 volume window (+-7 TRs) where the average timecourse for one participant’s parcel is correlated with the average timecourse of every other parcel across all remaining participants. The sliding window is calculated for every volume (step=1) to generate a Pearson correlation timeseries equivalent to the number of volumes in each movie.

Because our behavioral metrics span only the human annotated events and are not at the volume level, we selected the median volume in each annotated event, given that because of the sliding window the median volume would provide the most coverage of a specific event. The first and last 7 TRs of each movie were excluded from the analysis due to the fixed sliding window. Once the story-level dynamic intersubject functional correlation (dISFC) timecourses were the same dimension as our behavioral metrics, we calculated the Pearson correlation between each cognitive metric and these timecourses for each parcel-to-parcel comparison, generating 6,555 ROI comparisons including intersubject connectivity across the same parcel.

#### 2.8.3 Peak behavioral ISFC

With these ISFC values computed, we then used an analysis approach developed by Brandman et al. (2021) to compare the performance of Sequentiality with other automated and human-generated metrics of event properties. For each metric (Sequentiality, Causal Centrality, Semantic Centrality, Importance), we selected the five events with the highest values and then evaluated ISFC for those events across three networks of interest (default mode network (DMN), dorsal attention network (DAN), and ventral attention network (VAN)). ISFC values were averaged across the parcels in each network and calculated for each individual participant relative to the rest of the sample. Event onsets were shifted 3 TRs (4.5 seconds) to account for delays in the hemodynamic response function (HRF).

To ensure ISFC at peak Sequentiality events in the DMN was unique to the specific events selected, we computed statistical significance by comparing the mean ISFC across events for each participant with 1000 permutations of randomly selected events. This created a null distribution with the mean ISFC which allowed for estimating the p-value of the effect as the percentile of ISFC across this distribution. Our relationship of interest is how Sequentiality maps onto the ISFC in the DMN, so that is the only relationship we calculate significance with null permutations.

## 3. Results

We used our open-weight ensemble tool to re-analyze behavioral and neural data from three experiments. Specifically, we evaluated its flexible application across stimulus classes, correspondence to existing manual and automated measures, and ability to reproduce relevant behavioral and neural findings, now interpretable as reflecting the specifically computationally-defined quantity of *contextual information gain*. None of the reported analyses were preregistered, but each analysis is a direct parallel of those used in the studies employing manually generated ratings, specifically the univariate analyses match those from Lee & Chen (2022) and the intersubject functional analyses are from Brandman et al. (2021).

### 3.1 Relationship of Sequentiality to human- and computer-generated event properties

We first used the FilmFestival dataset (Lee & Chen, 2022; Section 2.7.1) to investigate the relationship between Sequentiality and existing quantifications of event properties. We computed two versions of the Sequentiality metric that differed in how informative the topic was with respect to the contents of each stimulus. Naive knowledge topics consisted of a concatenation of “A short story about” with the title of each movie. Outcome knowledge topics consisted of “A short story about” with a brief summary describing the plot of each movie (see Supplemental Materials for a full list of outcome knowledge topics).

To investigate the relationship between Sequentiality computed in these two ways, we computed the Pearson correlation between Sequentiality and each of the event property metrics included in the FilmFestival dataset; all quantities were z-scored within story before being entered into the correlation. We anticipated that the causal, not semantic structure of narratives would align with contextual information gain, and ran the correlations across the two metrics along with importance to see which features are particularly relevant to Sequentiality. As shown in Figure 2, both naive and outcome-knowledge Sequentiality was significantly correlated with human ratings of event importance (r_naive = 0.36, p_naive < .001; r_outcome = 0.23, p = .003) and causal centrality (r_naive = 0.24, p_naive = .003; r_clairvoyant = 0.27, p = .001). Critically, neither type of Sequentiality was associated with semantic similarity of events as computed using Universal Sentence Embeddings (USE; r_naive = 0.01, p_naive = .894; r_clairvoyant = 0.02, p = .783). To validate the difference between the relationship of causal as opposed to semantic centrality with Sequentiality we ran a fisher z-test between the two correlations and found they were significantly different (Z_naive=2.03, p_naive=.042; Z_clairvoyant=2.18, p_clairvoyant=.029),These findings thus demonstrate that Sequentiality tracks a psychologically relevant quantity that is not captured by existing automated methods for quantifying the dynamic properties of narrative stimuli, and is insensitive to the semantic fluency of sentences in a narrative.

**Figure 2:**
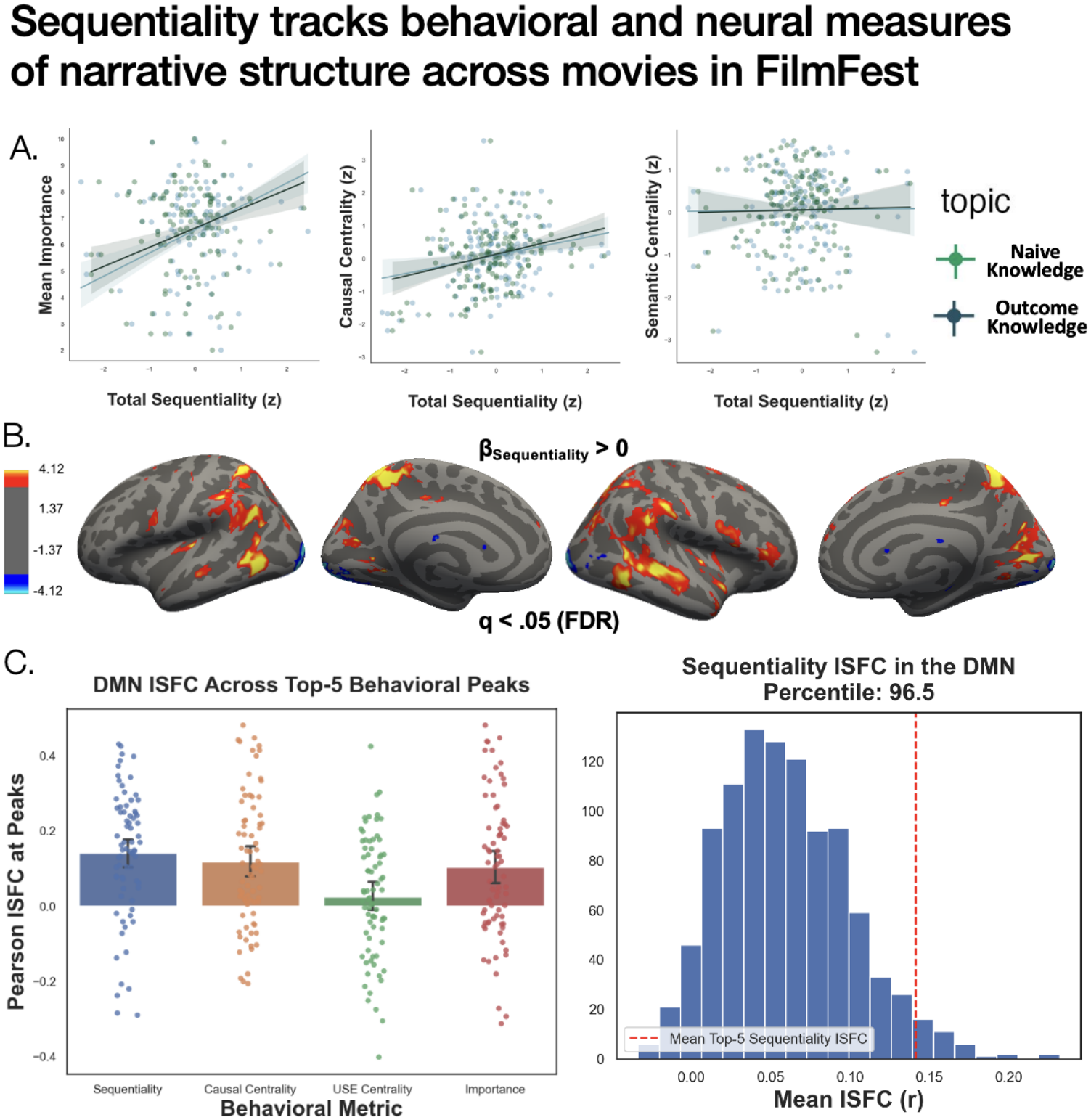
(A) Sequentiality is uniquely associated with human-generated ratings of event properties. Each point in the display corresponds to the z-scored Sequentiality value of each event/sentence. Shaded areas reflect 95% confidence intervals. (B) Whole brain univariate activation using Sequentiality as an event-related parametric modulator (*β*_Sequentiality_ > 0). Values were corrected using the False Detection Rate at q<.05 and cluster thresholded at 10 voxels. (C) Dynamic intersubject functional correlations (dISFC) at the median volume of each of the top-5 highest values of each behavioral metric. Each individual point is an individual participant’s ISFC at one of their five behavioral peaks (n=100). Following the original approach (Brandman et al., 2021), we compared ISFC at Sequentiality peaks to 1000 permutations of five random events, and plotted the percentile of peak ISFC.

### 3.2 Relationship of Sequentiality to neural activity during movie viewing

Given the reliable but moderate correlations with behavioral labels identified above, we next asked whether our metric corresponded to neural activity at the behaviorally-identified events. To do this, we turned to investigating how Sequentiality is related to BOLD activity measured during movie-watching in the FilmFestival dataset. First, we performed a whole-brain univariate activation analysis to identify the cortical regions whose BOLD activity scaled with Sequentiality (3.2.1). Then, we computed dynamic intersubject functional correlations (ISFC) to quantify the effect of Sequentiality on aligning neural activity across subjects during movie viewing.

#### 3.2.1 Univariate Analyses: Whole-brain Parametric Modulation by Sequentiality

As shown in Figure 2B, our univariate analyses revealed a diverse array of cortical regions whose activity was significantly modulated by Sequentiality. Positive associations were found for regions in the default mode, dorsal attention, and ventral attention networks, whereas negative associations were found for early occipital regions. Modulation of activity in the DMN is consistent with previous work demonstrating DMN modulation by the causal structure of FilmFest narratives (Lee & Chen, 2022; Figure S6).

#### 3.2.2 Intersubject Functional Connectivity at “Peak” Narrative Events

Given these univariate results, we focused our ISFC analyses on the default mode network along with dorsal and ventral attentional networks. The results for attentional networks are visualized in the Supplementary Materials. Following the approach developed by Brandman et al. (2021), we selected the moments in each narrative stimulus corresponding to the five highest values of each of our narrative metrics (Sequentiality, Causal Centrality, Importance, and Semantic Centrality) and computed the ISFC at the median volume of each event. Volume-wide timeseries were downsampled to behavioral timecourses, selecting the center ISFC values in each event similar to previous approaches (Brandman et al., 2021). To evaluate differences in cognitive measures across functional networks we ran a repeated measures ANOVA comparing the intersubject coactivation in parcels across the DMN, dorsal attention, and ventral attention networks with the previously listed cognitive measures. The ANOVA revealed a significant main effect of network (F(2,28)=50.58, p<.001, η□^2^ =.783) and notably a significant interaction between cognitive measure and functional network (F(6,84)=3.88, p=.002, η□^2^ =.217). Given our primary metric of interest is Sequentiality, we ran an additional repeated measures ANOVA with just Sequentiality and evaluated the relationship between network and ISFC, and found a significant main effect of network (F(2,28)=19.58, p<.001, η□^2^ =.583). Post-hoc contrasts revealed that moments of high Sequentiality were associated with ISFC values significantly greater than 0 in the DMN (t(14)=6.05, p<.001) and dorsal attention network (t(14)=7.91, p<.001), with no effect of Sequentiality on ISFC in the ventral attention network (t(14)=2.09, p=.166; Figure S4; all p-values Bonferroni corrected). We focused the remainder of our analyses on the DMN, given strong empirical evidence for its involvement in narrative stimulus processing (Ritchey et al., 2015; Barnett et al., 2020; Baldassano et al., 2018; Chen et al., 2017; Simony et al., 2016).

We next aimed to compare our automated metric of contextual information gain with previously calculated narrative metrics to determine its relative efficiency in capturing shared activation in the DMN. The left panel of Figure 2C demonstrates that moments with high Sequentiality were equally as effective at aligning neural activity in the DMN across subjects as human-generated ratings of event importance and causal centrality. Critically, events with high semantic centrality—the most prominent existing automated metric—did *not* significantly align cross-subject activity in this network. The right panel of Figure 2C further demonstrates that intersubject DMN correlations were significantly more aligned during high Sequentiality events than other events (1000 permutations, p=.035). These findings have two important implications. First, the DMN appears to be significantly associated with contextual information gain, as proxied by Sequentiality. Second, Sequentiality is a useful automated metric for studying narrative processing in the brain: it performs just as well as human ratings in aligning neural activity, and captures coactivation that existing automated metrics are not sensitive to. Taken together, these findings strongly motivate the use of Sequentiality as a tool for studying narrative processing in the brain and behavior and supporting the idea that this automated tool can scale up existing approaches.

### 3.3 Sequentiality captures dynamics within individual narratives

Having demonstrated that Sequentiality is associated with human ratings and neural activity as measured across all movies in the FilmFestival dataset, we next turned to investigating the ability of Sequentiality to capture canonical effects of narrative structure *within* single narratives. First, we show that Sequentiality reproduces effects of narrative “twists” on human ratings of both audiovisual and auditory-only stimuli (Section 3.3.1). Then, we show that dips in Sequentiality are significantly associated with human judgments of event boundaries in longer narratives (average duration > 7 minutes; Section 3.3.2). Together, these findings further support the utility of Sequentiality as an automated metric for quantifying dynamic and psychologically-relevant properties of narrative stimuli.

#### 3.3.1 Sequentiality reproduces effects of narrative “twists” on behavior

Narrative stimuli with “twists”—events that trigger a re-evaluation of the plot and/or characters in a story—are powerful tools for investigating dynamic reorganization of narrative encoding and its effects on subsequent memory (Sava-Segal et al., 2026; Zadbood et al., 2022). Sequentiality values are inherently conditioned on the perceived topic of a narrative (Figure 1A), and thus are well-suited to capture effects of prior knowledge of narrative “twists” on the contextual information gain of events occurring pre- and post-twist. We used two different stimuli with narrative twists to investigate this: an audiovisual narrative (*The Boyfriend*) from FilmFestival and an auditory-only narrative (*The Dark End of the Mall*) used in a recent study by Sava-Segal et al. (2026).

These narrative stimuli allow us to conduct a validity check on our re-interpretation of Sequentiality as a proxy of contextual information gain: how much information each observation adds to the dynamically-evolving history of observations comprising the “context” of narrative encoding. Because this quantity is computed relative to a user-defined “topic”—analagous to a human observer’s prior knowledge of the plot of the narrative—we ought to observe higher Sequentiality values for observations *after* a twist, and this difference should be amplified for topics reflecting “naive knowledge” relative to topics that contain information about the twist (“outcome knowledge”). For both analyses testing this prediction, we binned the standardized annotations into two categories: events occurring before vs. after the twist (“pre” vs “post” twist in Figure 3A).

**Figure 3:**
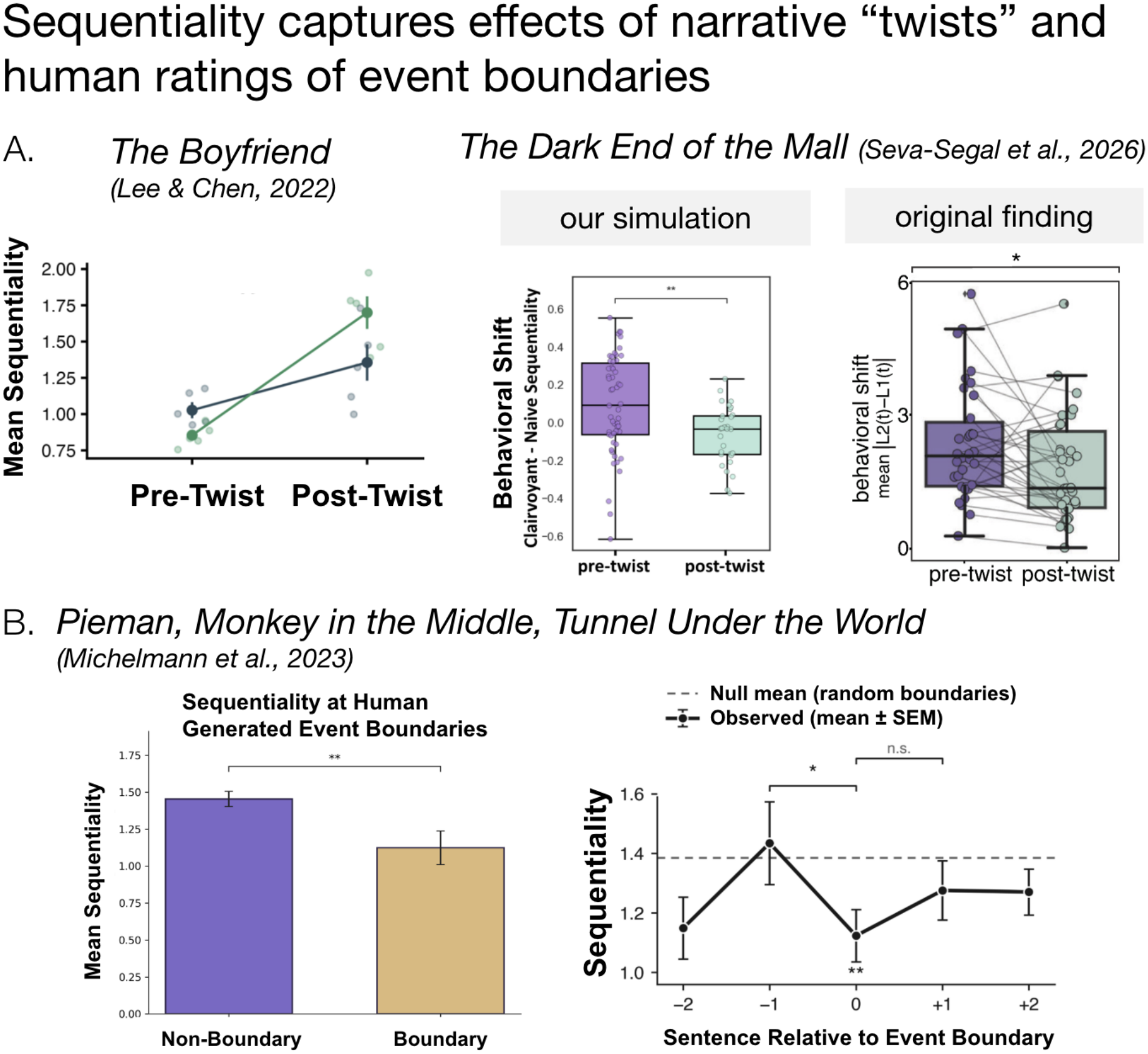
Sequentiality captures canonical structural effects within individual narratives. (A) Sequentiality is sensitive to “twists” within a single narrative. Changes in the outcome-information applied to the topic (*τ*) influence sequentiality differently before and after narrative-twists. Each point represents the ensemble Sequentiality for each sentence. *The Boyfriend* plot compares a naive and outcome-aware topic, where *The Repeat Viewing* plot uses the difference between outcome and naive Sequentiality values to mimic the influence of ‘repeat-viewing’ found in Seva-Segal et al., 2026. (B) Sequentiality dips at human-labeled event boundaries. They show this effect with respect to typical sentences (left) and the preceding sentence (right), and this result holds across multiple audio narratives. Permuted boundaries are shuffled event boundaries permuted 10,000 times; error bars are standard error.

As shown in the left panel of Figure 3A, Sequentiality values computed on standardized annotations of *The Boyfriend* exhibited precisely this pattern. If Sequentiality signals contextual information gain, then we should expect to see the values rise immediately after a twist for the topic that does not signal the outcome of that twist. Consistent with this interpretation, a linear model returned a significant interaction with sentence position (pre- vs. post-twist) and topic type, such that the average Sequentiality of events happening after the “twist” was significantly higher for the naive-knowledge topic relative to the outcome-knowledge topic (β = 0.5165, t_12_ = 4.306, p = .001). Before the twist, Sequentiality values computed with the naive-knowledge topic (light blue) were lower than the values generated with the outcome-knowledge topic (dark blue). After the twist, however, the naive-knowledge Sequentiality values were significantly greater. These findings thus directly support our contextual information gain interpretation of the Sequentiality metric and highlight the utility of user-defined topics as a *prior expectation* to the incoming information.

The findings displayed in the middle panel of Figure 3A show that manipulating the topic of Sequentiality and binning sentences into pre- and post-twist allows us to capture behavioral effects of multiple listens to a twist narrative as reported by Sava-Segal et al. (2026). The original study found that human ratings of character likability made *before* the narrative twist exhibited greater variability between listens than ratings made *after* the narrative twist (rightmost panel of Figure 3A), indicating a greater “shift” in behavior for pre- vs. post-twist events. This difference can be interpreted as reflecting effects of prior knowledge on the consistency of narrative evaluation: once observers know that a character they initially perceived as good is actually bad (i.e., after the twist in the first listen, or at any point in the second listen), their ratings of likeability decrease accordingly.

We simulate repeat viewing by computing Sequentiality with both naive- and outcome-knowledge topics, and then use the difference in these values as our proxy for changes in likability ratings. As in the original study, we found that changes in Sequentiality were greater for sentences occurring pre- vs. post-twist (t_94_=3.011, p=.003), reflecting human subject’s reevaluation—or differing evaluation—of likability as a function of prior knowledge of narrative twist. In addition to lending complementary support for our contextual information gain interpretation, these results further demonstrate the ability of the Sequentiality metric to capture psychologically-relevant properties of dynamically evolving narrative stimuli.

#### 3.3.2 Sequentiality tracks human-labeled event boundaries

For a final demonstration of the utility of Sequentiality for analyzing narratively-structured stimuli, as well as another test of our contextual information gain interpretation, we investigated how *dips* in Sequentiality relate to human judgments of event boundaries. If Sequentiality approximates how much information an observation adds to the temporally-evolving “context” of narrative encoding, then events with *low* Sequentiality can be interpreted as reflecting a shift or boundary in this continuous context window. To investigate this, we utilized existing data from a study that collected human ratings of event boundaries in auditory-only narratives (Michelmann et al., 2023; see Methods for more details). Because the narratives in this dataset exceeded the context window of some of our models, we calculated Sequentiality at four history lengths (2, 4, 6, and 8) and averaged each sentence’s Sequentiality value across them. The Sequentiality values were then averaged across all five models in the ensemble to produce one value per sentence in each story. Using the human ratings obtained by Michelmann et al. (2023), we then categorized each sentence as containing an event boundary or not, and binned the previously calculated Sequentiality values accordingly. A linear model revealed that Sequentiality values were lower for sentences containing human-judged event boundaries than for those without an event boundary (β = −0.33, t(527) = −2.65, p = .008), and that this effect was consistent across stories (|β| < .04, |t|(527) ≤ 0.41, p ≥ .68; Figure 3B).

To complement this finding, we next investigated how the *timecourse* of Sequentiality values related to human judgements of event boundaries (rightmost panel of Figure 3B). We obtained one timeseries of Sequentiality values per story, and created a null distribution by randomly placing the same number of boundaries within each story 10,000 times and computing the average Sequentiality for those stories. We then looked at Sequentiality values at and around sentences that were labeled as event boundaries by human subjects. We averaged a window of +/-2 sentences around each event boundary in all of the per-story time series. From a Monte Carlo permutation test, we found that sentences at event boundaries are lower than expected under the null distribution (one-sided, p=.005). Further, we see a statistically significant dip between the sentence right before the boundary and the sentence at the boundary when compared to the same sentence pair in the null (two-sided, p=.028). The difference between the Sequentiality values of the event boundary and the next sentence was not significant (two-sided, p=.331). Taken together, these findings strongly support both our re-interpretation of Sequentiality as a proxy of contextual information gain ***and*** the utility of this quantity for capturing psychologically-relevant dynamic properties of narrative stimuli.

Across three complementary analyses, we show that Sequentiality identifies dynamic narrative properties just as well as human subjects raters and better than existing similarity-based automated metrics, thus empirically demonstrating its utility for scaling up analyses in naturalistic neuroscience.

## 4. Discussion

We developed an automated, open-sourced metric representing contextual information gain throughout narrative events. This metric, Sequentiality, utilizes five open-weight LLMs that can be extended or replaced as desired. The model is conditioned on a user-generated topic, that can serve as its prior expectation, and then evaluates each upcoming sentence sequentially and compares it to its topic and a predefined sliding window of preceding sentences.

When we evaluated Sequentiality on several audiovisual narratives, we found that it is positively correlated with the importance and causal structure of narrative events, but notably not the semantic structure. This distinction between causal and semantic structure was not originally found across cognitive measures in the original dataset (Lee & Chen, 2022), and is a promising direction for future questions surrounding the properties of Sequentiality that effectively disambiguate the two metrics. When we evaluate stories with a narrative twist, we find running Sequentiality with outcome relevant topics lead to lower values compared to a naive prior. In line with our intuition, the more prior information the model has about the upcoming sentences, the lower it values the new information. We found this to be true not only when comparing the topics themselves, but also when comparing outcome and naive topics with intrasubject changes in character evaluations between first and repeat viewing of an audio narrative. The value of incoming information is expected to be shaped by internal goals, and we expect as these goals change, so would the degree of information gained in everyday experience. In line with this prediction, a promising future direction would be to adjust the metric to reflect a dynamic, mesoscale goal that is subsumed by the topic but can change throughout the narrative.

Neurally, Sequentiality is captured in subregions spanning the default mode network and both the dorsal and ventral attention networks. Previous metrics such as surprise (Brandman et al., 2021) and causal centrality (Lee & Chen, 2022) have shown a relationship between the DMN and cognitive metrics, and there has been an established relationship between attentional engagement in narratives and the neural activation in dorsal / visual streams of attention (Song & Rosenberg, 2023). Given there is no singular network implicated in contextual information gain, further research is needed to identify the importance of each selected subregion and how they might interact during narrative processing.

To effectively compare each metric across these functional networks, we utilized intersubject functional correlations between predefined parcels, and found a relationship between Sequentiality and DMN during the top five events with the highest contextual information gain. Evaluating inter-subject synchrony allows us to determine timepoints where a certain network was activating consistently across people. In other words, the DMN and Sequentiality coactivation implies events that included substantial, valuable information, were better represented and organized in more higher-order regions.

By definition, a high-Sequentiality event is one that adds topic-relevant information to the story. These ‘spikes’ reflect potential deviations or reevaluations to existing schemas. These changes in narrative information have been previously found in the DMN, and would provide further evidence that the network tracks changes in contextual information gain.

Alternatively, the ‘dips’ in Sequentiality during event boundaries could imply a variety of explanations. Previous research suggests that event boundaries occur when there is a substantial loss of information (Ashkenazi & Hart, 2026), while other research evaluates event boundaries under the framework of prediction errors (Rouhani et al., 2020) or alternatively working memory dynamics (Shim et al., 2024). The default mode network, specifically posterior medial regions, have been implicated in representing the higher-order representational timescales of continuous event structure (Baldassano et al., 2017). Future research is needed to disambiguate the reasoning behind these dips, and what properties of event structure they represent.

Our findings provide behavioral and neural support for a streamlined, automated system that reflects the influence of key computationally-defined quantities of information within narrative schemas. In doing so we provide an empirically grounded tool researchers can use to measure a ground truth level of informativeness throughout a narrative, guiding the expectation of subjective experience and advancing psychological theory. Validating this tool makes it feasible to scale naturalistic narrative science to larger corpora, such as those that extend across multiple episodes and which comprise heterogeneous stimuli across participants, now comparable using a common computational currency of contextual information gain.

