## Supplemental Materials for "Scaling up naturalistic narrative research: A reproducible large language model-based tool for evaluating contextual information gain"

### S1. Sensitivity of Sequentiality to sentence order in Hippocorpus

Although it has been shown that large language models can be relatively insensitive to word order permutations (Chen et al. 2024), the original paper describing the metric did not investigate whether Sequentiality displays the same insensitivity. To investigate this, we used a subset of Hippocorpus stories with the topic “my wedding,” shuffled each story 100 times, and then computed ensemble Sequentiality for the shuffled stories with history length $h \in \{1, 30\}$. We found that Sequentiality is indeed sensitive to the ordering of sentences within a narrative, such that the effect of story type (imagined, retold, or recalled) disappears when the sentences are shuffled (Figure S1). Additionally, a linear model confirmed that shuffling sentences significantly decreases Sequentiality values within each story type ($\beta = -0.3855$, $t5 = -12.468$, $p < .001$). Taken together, these findings validate the sensitivity of Sequentiality to the ordering of events in a narrative.

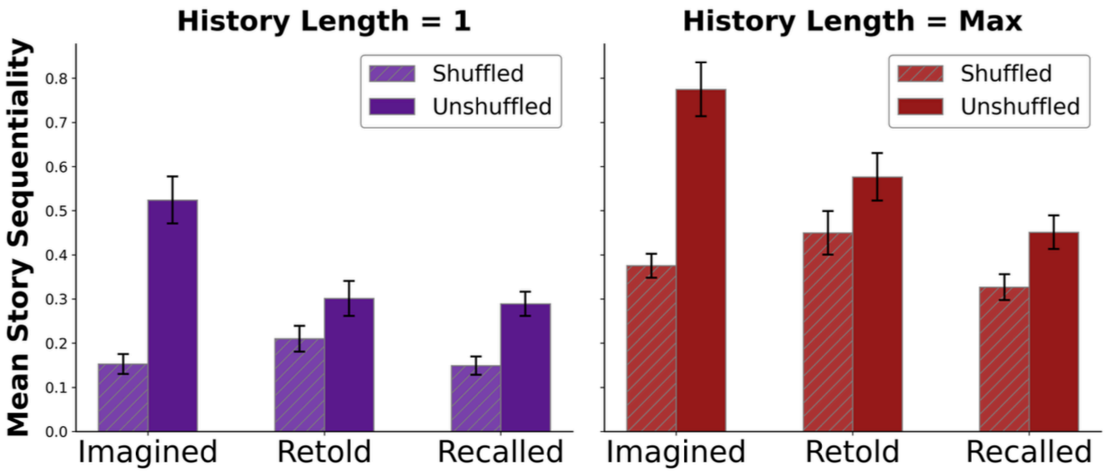


Figure S1: Sequentiality is sensitive to event sequences. Shuffling the order of sentences in Hippocorpus entries for the topic “My wedding” removes the effects of story type on average Sequentiality. Error bars depict standard error across models in the ensemble.

### S2. Details about fMRI acquisition and preprocessing

#### *fMRI acquisition*

fMRI scanning was conducted at Princeton Neuroscience Institute on a 3T Siemens Prisma scanner with a 64-channel head/neck coil (Lee & Chen, 2022). Functional images were acquired using a T2*-weighted multiband accelerated echo-planar imaging (EPI) sequence (TR = 1.5 s; TE = 39 ms; flip angle = 50°; acceleration factor = 4; shift = 3; 60 oblique axial slices;

grid size 96 × 96; voxel size 2 × 2 × 2 mm3). Fieldmap images were also acquired to correct for B0 magnetic field inhomogeneity (60 oblique axial slices; grid size 64 × 64; voxel size 3 × 3 × 2 mm3). Whole-brain high-resolution anatomical images were acquired using a T1-weighted MPRAGE pulse sequence. Scanning parameters for the anatomical images varied across participants (15 participants had 176 sagittal slices with voxel size 1 × 1 × 1 mm3; 6 participants had 192 sagittal slices with voxel size .9 × .86 × .86 mm3), as the anatomical images of a subset of participants were originally obtained for other projects unrelated to the current study.

#### Preprocessing of B0 inhomogeneity mappings

A B0 nonuniformity map (or fieldmap) was estimated from the phase-drift map(s) measure with two consecutive GRE (gradient-recalled echo) acquisitions. The corresponding phase-map(s) were phase-unwrapped with prelude (FSL 6.0.5.1:57b01774).

#### Anatomical data preprocessing

The T1-weighted (T1w) image was corrected for intensity non-uniformity (INU) with N4BiasFieldCorrection (Tustison et al. 2010), distributed with ANTs 2.3.3 (Avants et al. 2008, RRID:SCR_004757), and used as T1w-reference throughout the workflow. The T1w-reference was then skull-stripped with a Nipype implementation of the antsBrainExtraction.sh workflow (from ANTs), using OASIS30ANTs as target template. Brain tissue segmentation of cerebrospinal fluid (CSF), white-matter (WM) and gray-matter (GM) was performed on the brain-extracted T1w using fast (FSL 6.0.5.1:57b01774, RRID:SCR_002823, Zhang, Brady, and Smith 2001). Brain surfaces were reconstructed using recon-all (FreeSurfer 7.3.2, RRID:SCR_001847, Dale, Fischl, and Sereno 1999), and the brain mask estimated previously was refined with a custom variation of the method to reconcile ANTs-derived and FreeSurfer-derived segmentations of the cortical gray-matter of Mindboggle (RRID:SCR_002438, Klein et al. 2017). Grayordinate “dscalar” files (Glasser et al. 2013) containing 91k samples were also generated using the highest-resolution fsaverage as an intermediate standardized surface space. Volume-based spatial normalization to two standard spaces (MNI152NLin2009cAsym, MNI152NLin6Asym) was performed through nonlinear registration with antsRegistration (ANTs 2.3.3), using brain-extracted versions of both T1w reference and the T1w template. The following templates were selected for spatial normalization: ICBM 152 Nonlinear Asymmetrical template version 2009c [Fonov et al. (2009), RRID:SCR_008796; TemplateFlow ID: MNI152NLin2009cAsym], FSL’s MNI ICBM 152 non-linear 6th Generation Asymmetric Average Brain Stereotaxic Registration Model [Evans et al. (2012), RRID:SCR_002823; TemplateFlow ID: MNI152NLin6Asym].

#### Functional data preprocessing

For each of the 3 BOLD runs found per subject (across all tasks and sessions), the following preprocessing was performed. First, a reference volume and its skull-stripped version were generated using a custom methodology of fMRIPrep. Head-motion parameters with respect to the BOLD reference (transformation matrices, and six corresponding rotation and translation parameters) are estimated before any spatiotemporal filtering using mcflirt (FSL

6.0.5.1:57b01774, Jenkinson et al. 2002). The BOLD time-series (including slice-timing correction when applied) were resampled onto their original, native space by applying the transforms to correct for head-motion. These resampled BOLD time-series will be referred to as preprocessed BOLD in original space, or just preprocessed BOLD. The BOLD reference was then co-registered to the T1w reference using bbregister (FreeSurfer) which implements boundary-based registration (Greve and Fischl 2009). Co-registration was configured with six degrees of freedom. Several confounding time-series were calculated based on the preprocessed BOLD: framewise displacement (FD), DVARS and three region-wise global signals. FD was computed using two formulations following Power (absolute sum of relative motions, Power et al. (2014)) and Jenkinson (relative root mean square displacement between affines, Jenkinson et al. (2002)). FD and DVARS are calculated for each functional run, both using their implementations in Nipype (following the definitions by Power et al. 2014). The three global signals are extracted within the CSF, the WM, and the whole-brain masks. Additionally, a set of physiological regressors were extracted to allow for component-based noise correction (CompCor, Behzadi et al. 2007). Principal components are estimated after high-pass filtering the preprocessed BOLD time-series (using a discrete cosine filter with 128s cut-off) for the two CompCor variants: temporal (tCompCor) and anatomical (aCompCor). tCompCor components are then calculated from the top 2% variable voxels within the brain mask. For aCompCor, three probabilistic masks (CSF, WM and combined CSF+WM) are generated in anatomical space. The implementation differs from that of Behzadi et al. in that instead of eroding the masks by 2 pixels on BOLD space, a mask of pixels that likely contain a volume fraction of GM is subtracted from the aCompCor masks. This mask is obtained by dilating a GM mask extracted from the FreeSurfer's aseg segmentation, and it ensures components are not extracted from voxels containing a minimal fraction of GM. Finally, these masks are resampled into BOLD space and binarized by thresholding at 0.99 (as in the original implementation). Components are also calculated separately within the WM and CSF masks. For each CompCor decomposition, the k components with the largest singular values are retained, such that the retained components' time series are sufficient to explain 50 percent of variance across the nuisance mask (CSF, WM, combined, or temporal). The remaining components are dropped from consideration. The head-motion estimates calculated in the correction step were also placed within the corresponding confounds file. The confound time series derived from head motion estimates and global signals were expanded with the inclusion of temporal derivatives and quadratic terms for each (Satterthwaite et al. 2013). Frames that exceeded a threshold of 0.5 mm FD or 1.5 standardized DVARS were annotated as motion outliers. Additional nuisance timeseries are calculated by means of principal components analysis of the signal found within a thin band (crown) of voxels around the edge of the brain, as proposed by (Patriat, Reynolds, and Birn 2017). The BOLD time-series were resampled into standard space, generating a preprocessed BOLD run in MNI152NLin2009cAsym space. First, a reference volume and its skull-stripped version were generated using a custom methodology of fMRIPrep. The BOLD time-series were resampled onto the following surfaces (FreeSurfer reconstruction nomenclature): fsaverage5, fsaverage. Grayordinates files (Glasser et al. 2013) containing 91k samples were also generated using the highest-resolution fsaverage as intermediate standardized surface space. All resamplings can be performed with a single interpolation step by composing all the pertinent transformations (i.e. head-motion transform matrices, susceptibility distortion correction when available, and co-registrations to anatomical and output spaces). Gridded (volumetric) resamplings were performed using antsApplyTransforms (ANTs), configured with Lanczos interpolation to minimize the smoothing effects of other kernels (Lanczos 1964). Non-gridded (surface) resamplings were performed using mri_vol2surf (FreeSurfer).

### S3. Naive and Outcome Knowledge topics for FilmFestival stimuli

| Movie | Num Sent. | Naive Topic | Outcome Topic |
|---|---|---|---|
| Catch Me If You Can | 18 | A short story titled 'Catch me if you Can' | A short story about a famous con man being captured |
| The Record | 18 | A short story titled 'The Record' | A short story about a woman rapidly aging through her entire life |
| The Boyfriend | 18 | A short story titled 'The Boyfriend' | A short story about a robot woman replacing her robot boyfriend |
| The Shoe | 20 | A short story titled 'The Shoe' | A short story about a man and a woman mimicking each other until he is hit by a car |
| Keith Reynolds | 15 | A short story titled 'Keith Reynolds' | A short story about a man hoping for a promotion who ends up killing a coworker |
| The Rock | 19 | A short story titled 'The Rock' | A short story about a mountain becoming space debris |
| The Prisoner | 18 | A short story titled 'The Prisoner' | A short story about a man trapped in a strange village |
| The Black Hole | 18 | A short story titled 'The Black Hole' | A short story about a man getting trapped in a safe |
| Post-it Love | 16 | A short story titled 'Post-it Love' | A short story about two coworkers creating a mural of themselves |
| Bus Stop | 19 | A short story titled 'Bus Stop' | A short story about a man at a bus stop that turns into a dog |

### S4. Intersubject Functional Connectivity at “Peak” Narrative Events in Attentional Networks

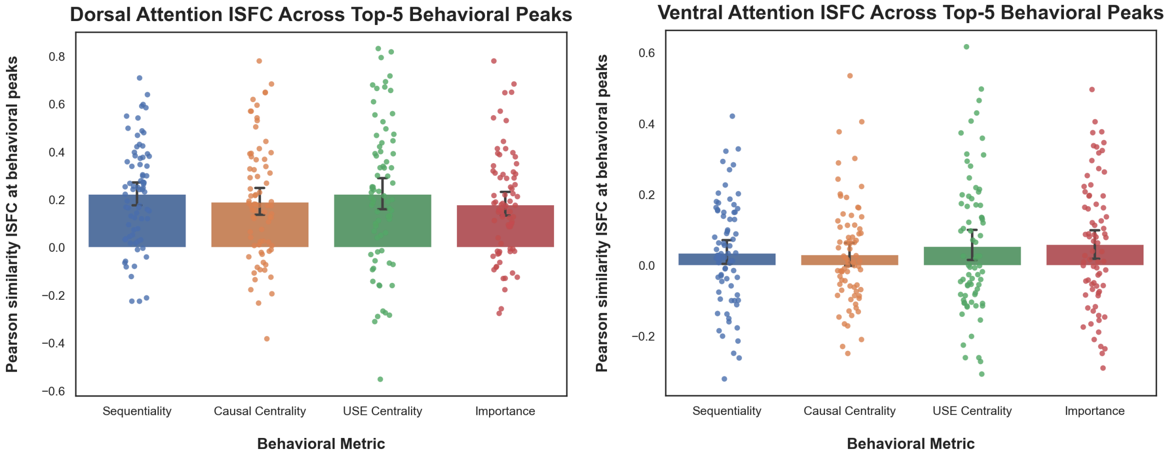


### S.5 Prompt used to standardize each narrative with an LLM

SYSTEM:
You convert narrative text into a condensed, standardized form while preserving a mapping back to the original sentences.

INPUT: A JSON array of pre-indexed sentences from a story or transcript:
[{"id": 0, "text": "..."}, {"id": 1, "text": "..."}, ...]

TASK:
Rewrite the narrative as a sequence of condensed sentences that together preserve all plot-relevant information (events, actions, states, causal relationships). For each condensed sentence, record which original sentence id(s) it draws information from. Do not attempt to identify or segment discrete "events" — condensation should follow naturally from combining related, adjacent information, not from imposing event boundaries.

CONSTRAINTS on every condensed sentence:
- Active voice only (no passive constructions).
- Strict Subject-Verb-Object word order.
- Neutral, factual tone — no dialogue, no quoted speech, no editorializing, no figurative language.
- Convert all dialogue into indirect narration of what was said, decided, or revealed (e.g. "Elena told him she was leaving" not "'I'm leaving,' Elena said").
- Each sentence should express one clear action, state, or relationship — combine source sentences only when their information is closely related and combining does not overload the sentence.
- Do not invent information not present in or directly implied by the source sentences.
- Do not omit plot-relevant information: named entities, causal links, and outcomes must be traceable to the condensed output.

- Preserve chronological order as it occurs in the source.

COVERAGE:
- Every input sentence id must be referenced by at least one condensed sentence's source_ids.
- A condensed sentence may draw from multiple source ids, contiguous or not, if their information is directly related.
- Do not skip source ids, even if they seem minor (e.g. brief dialogue tags, transitions) — fold their information into the nearest relevant condensed sentence.

OUTPUT FORMAT:
Return only valid JSON, no other text, in this exact structure:
{
  "condensed": [
    {"condensed_id": 0, "text": "...", "source_ids": [0, 1]},
    {"condensed_id": 1, "text": "...", "source_ids": [2]},
    ...
  ]
}

Do not include markdown code fences. Do not include explanation before or after the JSON.

USER:
{{PRE_INDEXED_SENTENCES_JSON}}

#### S.6 Comparing Univariate Activation with Sequentiality and original Causal Centrality findings (Lee & Chen, 2022)

Sequentiality vs Causal Centrality (Lee & Chen, 2022) (p<.001, uncorrected)

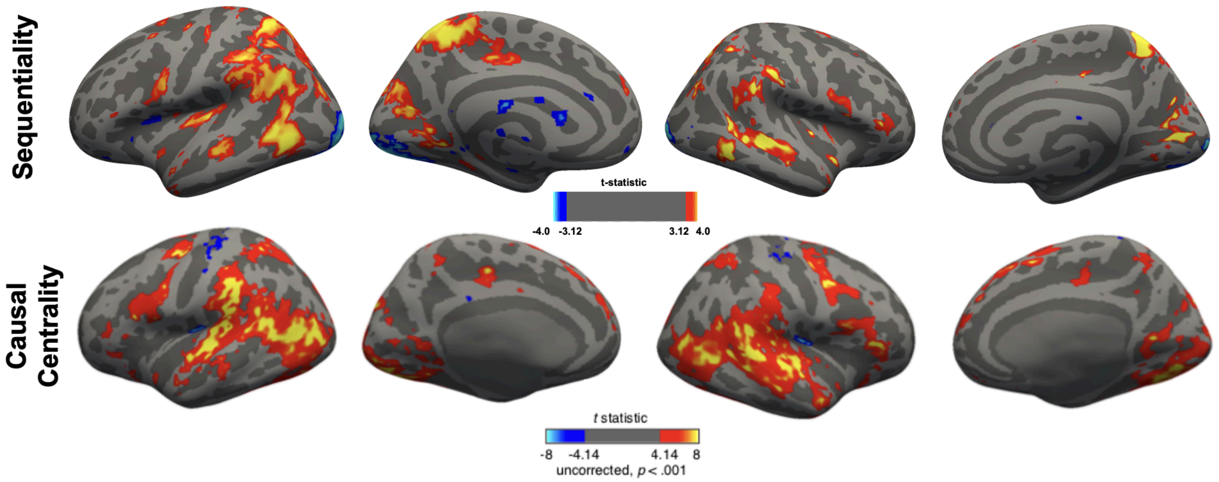
